# A Model for Resource Competition in CRISPR-Mediated Gene Repression

**DOI:** 10.1101/266015

**Authors:** Pin-Yi Chen, Yili Qian, Domitilla Del Vecchio

## Abstract

CRISPR-mediated gene regulation is known for its ability to control multiple targets simultaneously due to its modular nature: the same dCas9 effector can target different genes simply by changing the associated gRNA. However, multiplexing requires the sharing of limited amounts of dCas9 proteins among multiple gRNAs, leading to resource competition. In turn, competition between gRNAs for the same resource may hamper network function. In this work, we develop a general model that takes into account the sharing of a limited amount of dCas9 protein for arbitrary CRISPR-mediated gene repression networks. We demonstrate that, as a result of resource competition, hidden interactions appear, that modify the intended network regulations. As a case study, we analyze the effects of these hidden interactions in repression cascades. In particular, we illustrate that perfect adaptation to resource fluctuations can be achieved in cascades with an even number of repressors. In contrast, cascades with an odd number of repressors are substantially impacted by resource competition.

## I. INTRODUCTION

Synthetic genetic circuits have shown their potential in a number of applications, from energy, to environment, to medicine [1]. For complex and scalable synthetic circuits, effective and programmable synthetic transcription factors are the building blocks. CRISPR-Cas9 systems, one of the most exciting recent discoveries in biology, provide a simple and versatile tool for genetic modifications in various cell types and have been recently repurposed for transcriptional regulation [2]. Compared with other major classes of programmable synthetic transcription factors, such as ZFs [3], [4] and TALEs [5], CRISPR-mediated gene regulation offers unprecedented ease in multiplexing, which is vital for building complex gene circuits. The modular nature of this RNA-guided DNA recognition platform, where a single protein dCas9 can target different genes by changing the associated gRNA, makes RNA-guided transcriptional regulation precise, scalable and robust.

Repression through CRISPR interface (CRISPRi) and activation through CRISPR-mediated gene activation (CRISPRa) have been achieved in diverse organisms, including bacterial and eukaryotic cells [6]. For CRISPRi in bacterial cells, by pairing dCas9 with sequence-specific sgRNAs, dCas9-sgRNA complexes can efficiently inhibit the transcription of targeted genes. In mammalian cells, CRISPRi can be enhanced by fusing dCas9 to a transcriptional repressor domain, such as KRAB [7] and SID4X [8]. In addition to CRISPRi, CRISPRa has been created by fusing dCas9 to *ω*-subunit [9] of RNA polymerase in bacteria. The fusion of transcriptional activators, such as VP64 and p65AD [7], to dCas9 leads to activation in mammalian cells. Simultaneous activation and repression of genes was also established by using scaffold RNAs (scRNAs) [10]. The scRNAs encode information both for DNA target recognition and for recruiting a specific repressor or activator protein. By sharing the same dCas9 protein, dCas9-scRNAs can simultaneously repress or activate multiple genes in the same cell.

Previous work on building CRISPR-based circuits has demonstrated the potential of using CRISPR-mediated gene regulation to build layered, complex and scalable synthetic regulatory circuits [11]–[13]. However, as circuits become larger, a greater number of gRNAs are expressed, and thereby, competition for a finite pool of dCas9 may cause unintended interactions. In addition, since high levels of dCas9 concentration are toxic, leading to reduced cell growth [11], one cannot mitigate competition by arbitrarily increasing dCas9 production. Therefore, it is important to determine how competition for a finite amount of dCas9 affects the emergent circuit behavior.

The effects of resource competition have been extensively studied for different cellular resources such as ribosomes [14] [15] and proteases [16] [17]. These studies demonstrated significant effects of resource competition and provided model-guided methodologies to minimize the resulting effects. However, to the best of our knowledge, there has not been any study about the effects of competition for dCas9 in CRISPR-based genetic circuits.

In this paper, we develop a simple ODE model whose state variables are the concentrations of the gRNAs and output proteins. The model, which explicitly accounts for the effects of dCas9 sharing, is general enough to capture arbitrary CRISPRi networks in both bacterial and mammalian cells. The steady state I/O responses are discussed in parallel networks. In addition, the “hidden” interactions, which are all activations for CRISPRi-based circuits, are added to the interaction graph of the system. Finally, the model is applied to CRISPR-based repression cascades and illustrates that even-stage cascades have better adaptation to the competition effect than odd-stage cascades under certain assumptions.

The rest of the paper is organized as follows. In Section II, we introduce the modeling framework for parallel repression networks and the concept of I/O response. In Section III, we introduce the general modeling framework, which is applicable to any repression-only network, such as repression cascades. Then, the concept of competition-induced hidden interaction is defined. In Section IV, an n-stage repression cascade example is detailed as an application of our general modeling framework. The design guideline for perfect adaptation to resource competition is provided.

## II. RESOURCE COMPETITION IN PARALLEL NETWORKS

### A. Modeling Framework

There is a wide variety of CRISPR-based platforms for gene regulation. Both gRNA and dCas9 can be modified for a particular scenario, including repression or activation in bacterial or mammalian cells. In either case, there is always a pool of dCas9 or modified dCas9, such as dCas9-*ω* or dCas9-VP64, shared by sequence-specific gRNAs or scRNAs. Therefore, in this model, we neglect the conformational detail of each component, and lump the key species into two: the resource (dCas9 or modified dCas9) and the users (gRNAs or scRNAs), by naming them d and g, respectively. As shown in Fig. 1, we first consider a network with *n* users without regulatory interactions among them (i.e., parallel network). For CRISPR-mediated gene regulation, d pairs with *g_i_* (where *i* = 1, 2,…, *n*), forming dCas9-gRNA complexes (c_i_), which act as transcription factors. These transcription factors then interfere with the transcription elongation of the target genes (D_i_), to form complexes (C_i_):

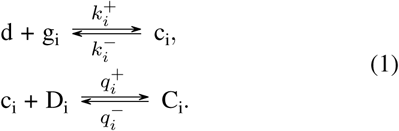

**Fig. 1.**
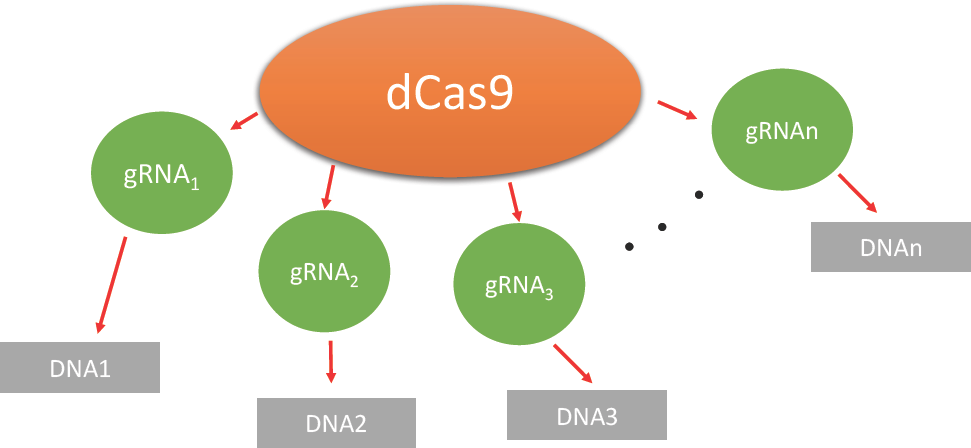
dCas9 is the shared resource among multiple gRNAs.

For CRISPR-mediated gene repression, termed CRISPRi, the free target genes (D_i_) are transcribed and then translated into output proteins (Y_i_). The concentrations of output proteins (*Y_i_*) are used to study the repression level:

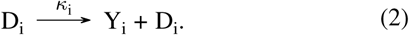

For CRISPR-mediated gene activation, termed CRISPRa, the transcriptionally active complexes (C_i_) produce the output proteins (Y_i_):

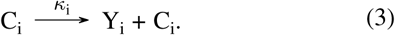

In activation and repression, the sequestration mechanism for resources is identical (1). For simplicity, we focus on CRISPR-mediated gene repression for the remainder of this paper. The analysis of activation can be dealt with similarly and is left for future work. In our model, the decay rate of the complexes are neglected since we assume that *k^−^_i_* and *q^−^_i_* are much greater than the decay rate of complexes. The production rates of gRNA *i* are *u_i_*. The decay (i.e., dilution plus degradation) rates of gRNAs and output proteins are and, respectively, which we assume to be constant for all nodes without loss of generality. We use *d*, *g_i_*, *c_i_*, *C_i_*, *D_i_* and *Y_i_* to represent concentrations of species d, g_i_, c_i_, C_i_, D_i_ and Y_i_, respectively. Consequently, based on chemical reactions (1) - (3), we have the following ODE model from mass action kinetics:

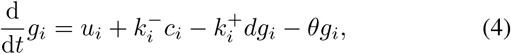

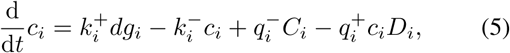

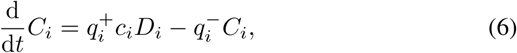

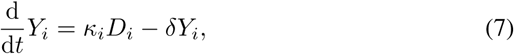

where indices *i* = 1, 2,…, *n*. Since the total concentrations of dCas9 and DNAs are conserved [18], we have

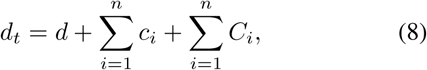

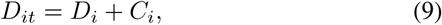

where *d_t_* and *D_it_* are the total concentrations of d and D_i_, respectively. Since the binding reactions in (1) are much faster than the production and decay of proteins and RNAs [18], the time derivatives in equations (5) and (6) are set to zero (quasi-steady state assumption). The complex concentrations in each node *i* at quasi-steady state (QSS) are as follows:

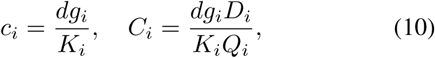

where dissociation constants *K_i_* and *Q_i_* are defined as:

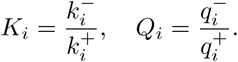

From (9) and (10), free DNA concentration (*D_i_*) at steady state can be written as:

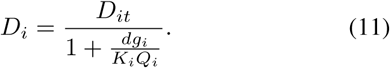

To obtain the free dCas9 concentration *d*, we substitute (10) and (11) into equation (8) to obtain:

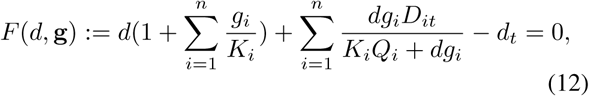

where we use vector g: = [*g*_1_,…, *g_n_*]^*T*^ to represent all gRNA concentrations in the system. For any given g, since *F* (g, *d*) is monotonically increasing with *d* for all *d* > 0 and ranges (−*d_t_*, +∞), equation (12) has a unique positive solution *d* =*d* (g) *>* 0. By substituting *c_i_* concentrations from (10) into (4), and free DNA concentration (11) into (7), the dynamics of (4)-(7) can be simplified to:

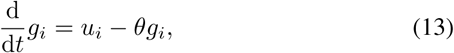

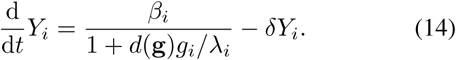

where

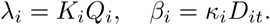

The above equations show that gRNA’s dynamics (13) depend only on its input. Instead of in (13), the competition effect is captured in (14), where resource availability *d* (g) appears in the denominator. For a constant input *u_i_*, steady state gRNA concentration 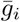 and output protein concentrations 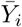 can be computed from (13)-(14):

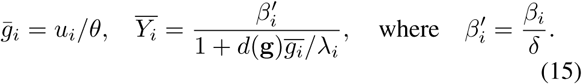

When there are abundant resources, dCas9 concentration remains approximately constant, d ≈ *d_t_*. The output concentration at steady state *Y_i_* can be written as:

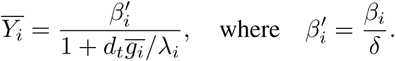

We perform numerical simulations on a two-gRNA parallel network based on the model we derived as an illustrative example (Fig. 3). As the production rate (*α_d_*) of dCas9 decreases, the regulation effect is altered. In Fig. 3, when the production rate of the resource decreases by 10 fold, the repression level diminishes by approximately 6 fold (the red line in the left figure), which is caused by the “hidden activation” (the red line in the right figure). In the right figure of Fig. 3, the input *u*_2_ is essentially uncoupled with *Y*_1_ when there are sufficient resources (the blue line). When the production of resource decreases by 10 fold, the “hidden activation” from *u*_2_ to *Y*_1_ becomes appreciable (about 5 fold). The diminished repression level and hidden activation level depend on not only the amount of gRNAs (*u*_1_, *u*_2_) and dCas9 (*_d_*), but also other parameters, such as dissociation constants (*K*_1_, *K*_2_, *Q*_1_, *Q*_2_), dilution rates (*θ*, *δ*), the amount of targets (*D_1t_*, *D_2t_*), and the output promoter strengths (*κ*_1_). The parameters used in the simulation are estimated from literature and preliminary experimental results.

**Fig. 2.**
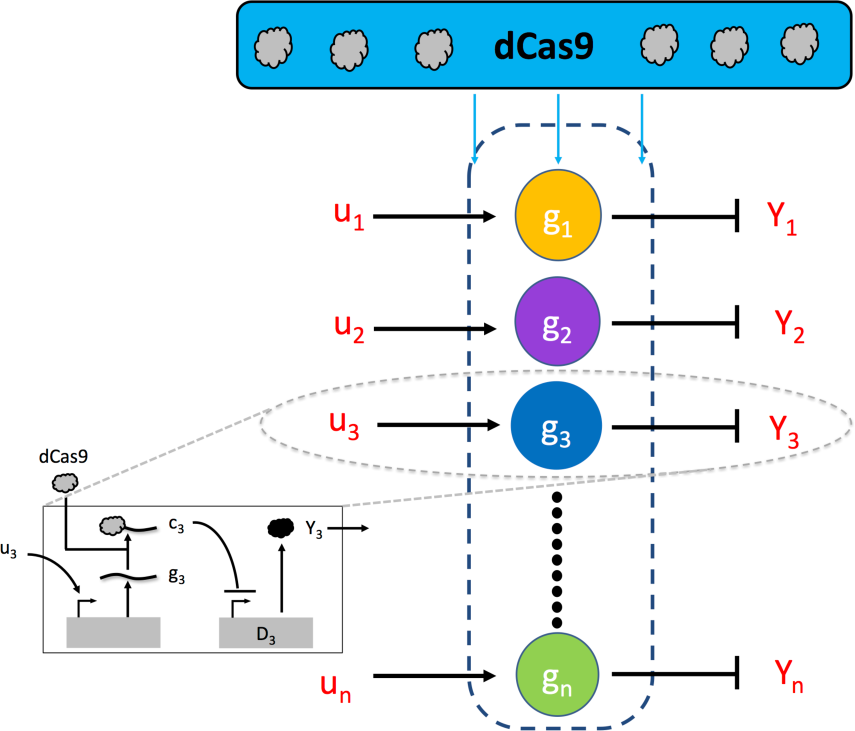
A parallel network with n gRNAs

**Fig. 3.**
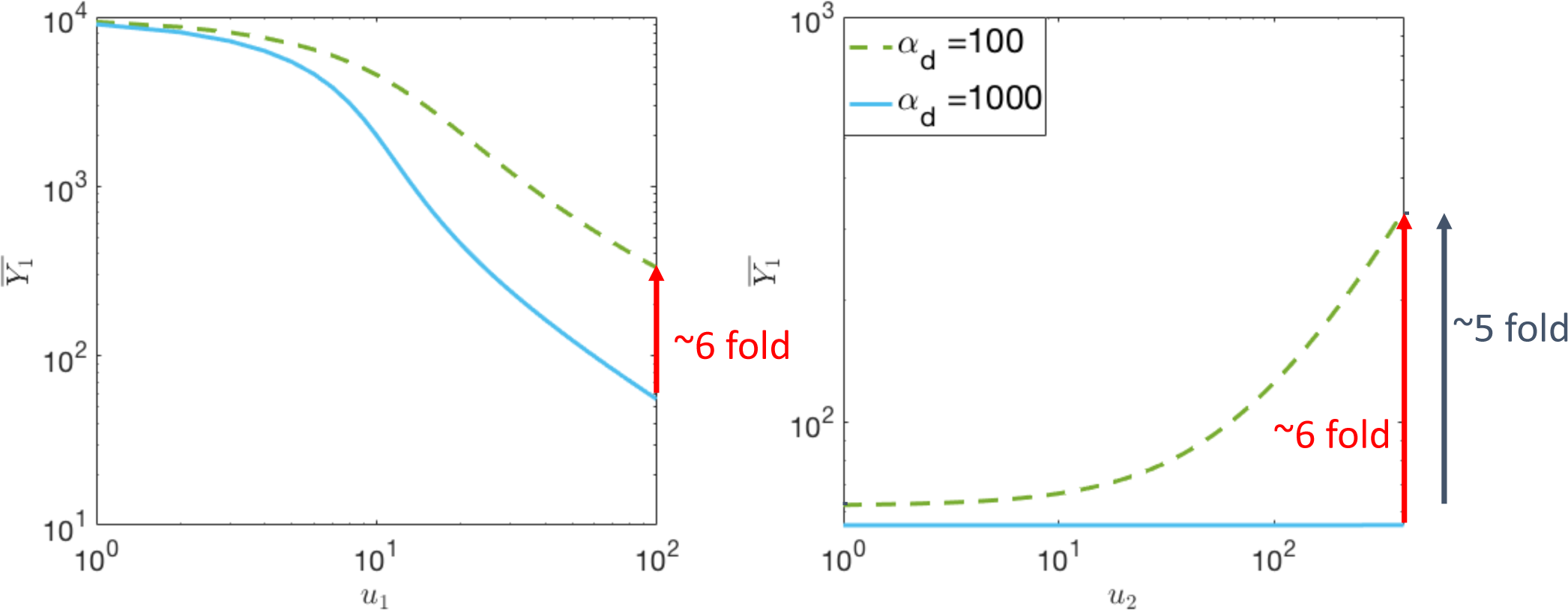
I/O steady state response of a parallel network with two gRNAs. The red lines in both figures represent the fold change of 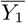 concentration from the case where the resource are abundant (*α_d_* = 1000) to limited (*α_d_* = 100) when *u*_1_ = 100 and *u*_2_ = 400. The black line in the right figure represents the fold change (”hidden activation”) from *u*_2_ = 0 to *u*_2_ = 400 when *_d_* = 100. In the left figure, *u*_2_ concentration is fixed at 400nMhr^−1^. In the right figure, *u*_1_ concentration is fixed at 100nMhr^−1^. Other Parameters: *D_1t_* =*D_2t_* = 10nM, *K*_1_ =*K*_2_ = 0.01nM [19], *Q*_1_ =*Q*_2_ = 0.5nM [20], *δ* = 1hr^−1^, = 100hr^−1^, *κ* = 1000hr^−1^.

### B. Conservation of dCas9

In the following section, the relation between the free amount of dCas9 (*d*) and gRNA concentrations (g) is studied. First, we consider two extreme scenarios to explicitly represent *d* as a function of g. Then, we investigate how *d* changes qualitatively with g, namely, *∂d*/*∂g_i_*. This information will be utilized later when we evaluate the sign of the I/O response and the interaction graph.

#### Senario (i): High Repression Level

When there are sufficient dCas9 (*d_t_*) and gRNAs (*g_ti_*) in the system, abundant repressors are formed. Under this scenario, all the targets are repressed, namely, the amount of repressed targets approximately equal to the total amount of targets: *C_i_* ≈ *D_it_*. Equation (8) can be written as:

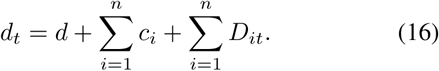

By substituting equation (10) into the above equation, we obtain the free dCas9 concentration:

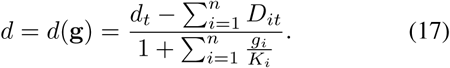

#### Senario (ii): Low Repression Level

When there are limited dCas9 protien or of gRNAs, most targets are not being repressed. Under this scenario, the amounts of free DNAs approximately equal to the total amounts of DNAs: *D_i_* ≈ *D_it_*. By substituting (10) into equation (8), we obtain:

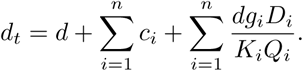

By substituting equation (10) into the above equation, we obtain the free dCas9 concentration:

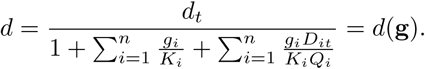

Next, we investigate the sign of *∂d*/*∂g_i_*. Since *∂F*/*∂d* ≠ = 0 for all *d* > 0 in (12), by the implicit function theorem [21], we have:

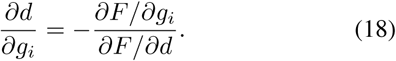

We explicitly find the sign of *∂F*/*∂g_i_* and *∂F*/*∂d*:

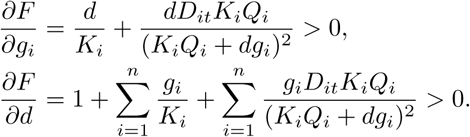

Consequently, according to (18), the sign of *∂d*/*∂g_i_* is guaranteed to be negative (i.e., sign[*∂d*/*∂g_i_*] < 0 for all *i*).

### C. Steady State I/O Response

When the resource is limited, the steady state I/O response reflects the competition effect. The following equation is the steady state I/O response from any input *u_j_* to any output 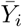:

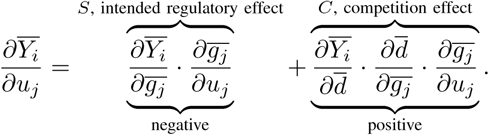

The first term (*S*) is due to the intended regulation (i.e. repression), and the second term (*C*) arises from unintended interactions due to resource competition.

#### Claim 1

The steady state I/O response of the parallel network in (15) satisfies:

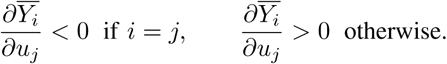

*Proof:* From equation (15), when *i* = *j*, we have:

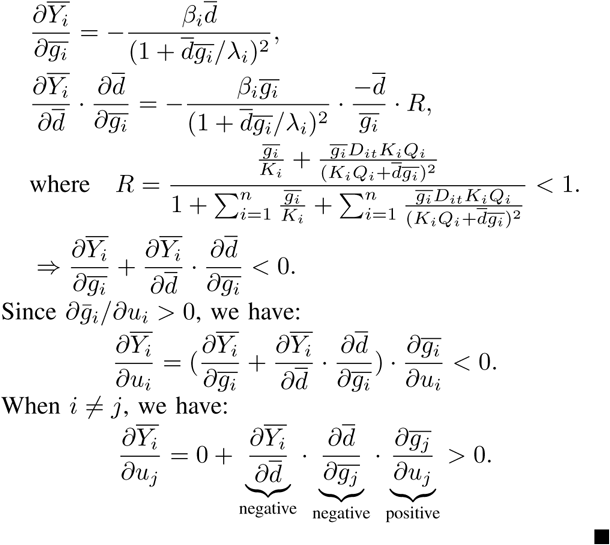

**TABLE I.** THREE TYPES OF NODES IN A CRISPRI GENE REGULATION CIRCUIT

| type | 1 | 2 | 3 |
| --- | --- | --- | --- |
| input | external inducer | dCas9-gRNA | dCas9-gRNA |
| output | dCas9-gRNA | dCas9-gRNA | protein |

#### Remark 1

The claim above implies that while resource competition and intended regulation always has opposite qualitative effect, the resource competition effect is always weaker when *i* = *j*. When *i* ≠ = *j*, there is no intended interaction, and the resource competition always causes hidden activation.

## III. RESOURCE COMPETITION IN GENERAL NETWORKS

In the previous section, we focus on a less general scenario where the repressors are restricted to target output proteins. However, complex CRISPR repression networks may have direct regulations between gRNAs (i.e. the repressors fromed by one gRNA and dCas9 repress the production of another gRNA in the network.) Therefore, in this section, we provide a general modeling framework that captures arbitrary CRISPR repression network. Here, we first give a high-level description of a CRISPRi gene regulation network in Section III-A, which includes a classification of 3 types of node in the network. We then describe the dynamics in each type of nodes (Section III-B) and introduce the resource constraint imposed on the network (Section III-C). We summarize the network resource competition model in Section III-D. We demonstrate in Section III-E how resource competition gives rise to hidden interactions among nodes.

### A. Preliminaries

A CRISPRi gene regulation circuit can be modeled as a network of *m* nodes. Depending on the biomolecular species each node takes as inputs and produces as output, it can be classified into one of the 3 types in Table I (see also Figure 4). We use index sets 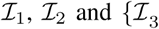 to denote the set of nodes that fall into type 1,2 and 3, respectively, and define 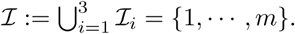 In a type 1 (2) node, an external inducer (a set of dCas9-gRNA complexes) regulates the transcription of a gRNA, which can bind with dCas9 to form a dCas9-gRNA complex as an output. In contrast, a type 3 node is regulated by a set of dCas9-gRNA complexes to produce a protein as output. We use 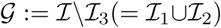 to represent the set of nodes that transcribes a gRNA. We let 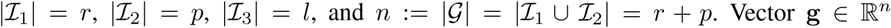 represent the concatenation of gRNA concentrations in these nodes, arranged according to the ascending order of their indices. The indexing starts with *r* nodes in 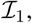 followed by *p* nodes in 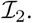 We use 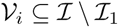 to represent the set of nodes regulated by node *i* (i.e., *targets* of node *i*). Similarly, we use 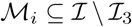 to represent the set of nodes that regulate the transcription in node *i* (i.e., *parents* of node *i*). We use vector 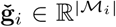 to represent the concatenation of gRNA concentrations in 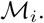 An example network with *m* = 6 nodes is shown in Figure 4. In this example, nodes of different types are filled with different colors. Specifically, we have 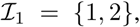 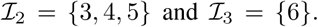 As an illustrative example, the set of parents to node 4 is 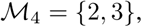 and therefore, 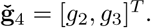 The set of targets of node 3 is *V*_3_ = {4, 5}.

**Fig. 4.**
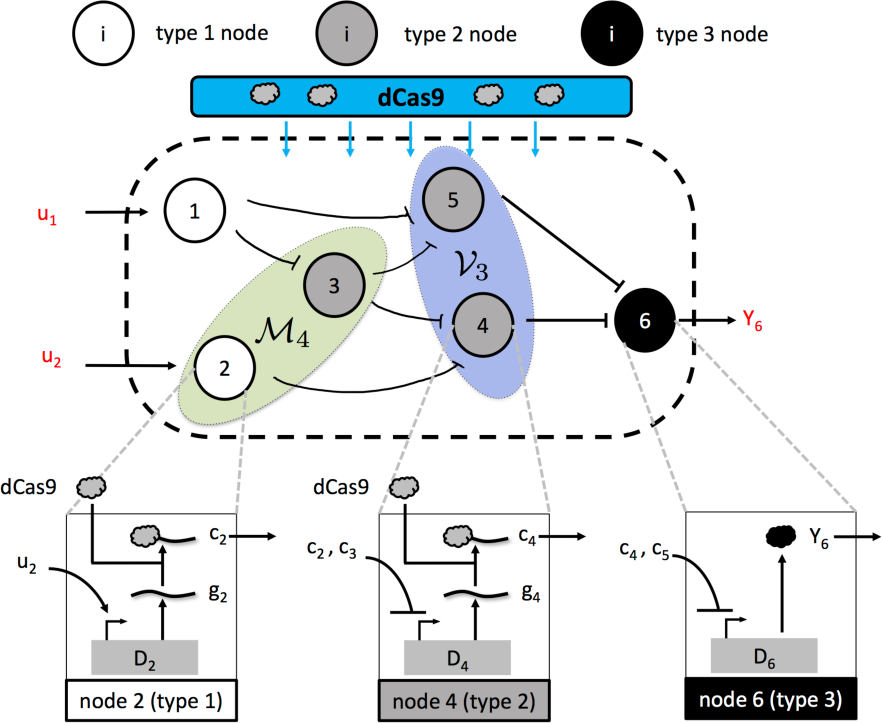
An example CRISPRi regulation network. This network can be modeled by the general modeling framework. It is composed of 6 nodes. Nodes of type 1, 2 and 3 are colored white, gray and black respectively. Biomolecular processes for node 2, 4 and 6 are shown as examples. The set of parent nodes to node 4 is 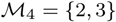 and the set of target nodes of node 3 is *V*_3_ = {4, 5}.

In what follows, we describe in detail the chemical reactions and dynamics in each type of node.

### B. Dynamics in a node

*1) <u>Type 1 node:</u>* A type 1 node 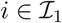 takes an external inducer u_i_ as input to produce a gRNA g_i_, which binds with free dCas9 (d) in the network to form an active regulatory complex c_i_ as output. These chemical reactions can be written as:

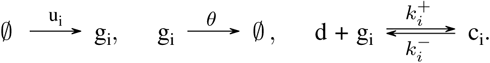

The output complex c_i_ can then bind with DNA of its target node *k* ∈ *V_i_*, D_*k*_, to repress its transcription:

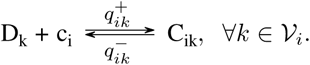

Based on the above chemical reactions, the dynamics of *g_i_* and *c_i_* in node *i* follow:

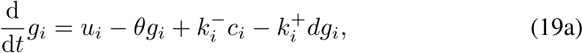

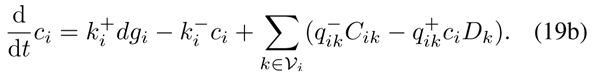

*2) <u>Type 2 node</u>:* The transcription of gRNA g_*i*_ from its DNA 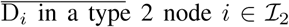 is repressed by a set of dCas9-gRNA complexes produced by its parents, resulting in the following chemical reactions for each 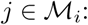

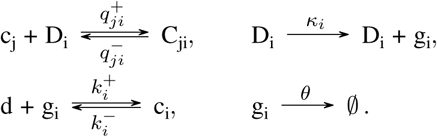

The output c_i_ can then repress a target node *k* ∈ *V_i_* by binding to its DNA *D_k_* to block transcription:

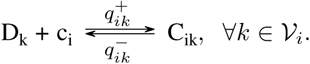

Based on these reactions, the dynamics of *C_ji_*, *g_i_* and *c_i_* can be written as:

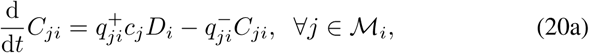

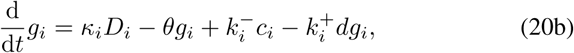

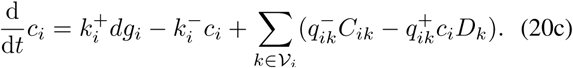

*3) <u>Type 3 node</u>:* A node 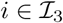 is repressed by its parent nodes to express a protein *Y_i_* from its DNA *D_i_* as output. In particular, dCas9-gRNA complexes *c_j_* produced by any node 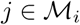 can block the transcription of Y_*i*_. We assume that when production of Y_*i*_ is not repressed, it takes place with a constant rate constant *κ_i_*. These processes can be described by the following chemical reactions:

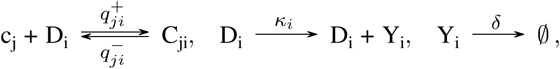

where *δ* is the dilution rate constant proportional to the specific growth rate of the host cell. These reactions can be described by the following ODEs:

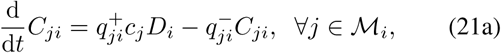

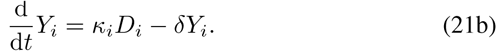

Now that we have studied the dynamics in all types of nodes, we are ready to factor resource competition into the model where they are connected.

### C. Conservation of dCas9

We assume that the circuit produces a limited amount of dCas9 (*d_t_*) available to all nodes, and therefore follows the conservation law:

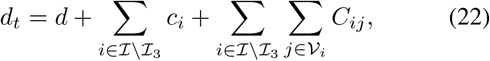

where we have summed up dCas9 1) bound to form DNA-dCas9-gRNA complexes (C*_ij_*), which only appears in type 2 and 3 nodes, and 2) bound to form dCas9-gRNA complexes *c_i_*, which only appears in type 1 and 2 nodes.

Assuming that binding reactions are much faster than the transcription, we can compute the QSS concentrations of *C_ij_* and *c_i_*. Specifically, by setting the time derivatives in equations (20a) and (21a) to 0, we find that

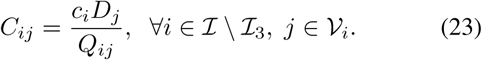

Note that in (19b) and (20c), for any 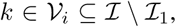 we must have 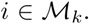. Therefore, we can substitute the result in (23) into (19b) and (20c), and find *c_i_* at QSS by setting the time derivatives to 0 in (19a) and (19b) to obtain

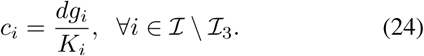

Substituting the results in (23) and (24) into the conservation law (22), we obtain:

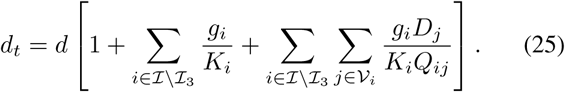

To find the free DNA concentration *D_i_* in (25), we assume that the total concentration of DNA *D_it_* in any node 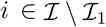 is conserved [22]:

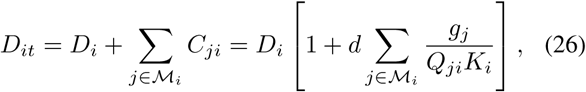

where we have substituted in the results in equations (23) and (24). To find free dCas9 amount, we use equations (25) and (26) then obtain:

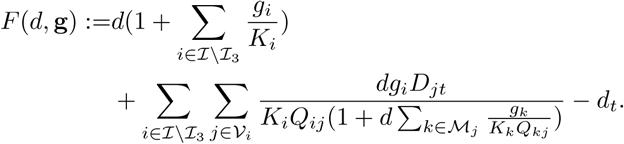

For any given g, since *F* (g, *d*) is monotonically increasing with *d* for all *d* > 0 and ranges (−*d_t_*, +∞), equation (12) has a unique positive solution *d* =*d*(g) > 0. In the following section, the relation between the free amount of dCas9 (*d*) and gRNA concentrations (g) is studied similarly to that of like the parallel network in section II. First, we consider two extreme scenarios. Then, we investigate *∂d*/*∂g_i_* qualitatively.

#### Senario (i): High Repression Level

When there are sufficient dCas9 (*d_t_*) and gRNAs (*g_ti_*) in the system, abundant repressors are formed. Under this scenario, all the targets are repressed, namely, the number of repressed targets approximately equals to the total number of targets: 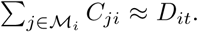 Equation (22) can be written as:

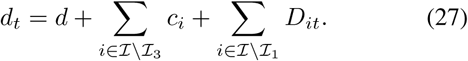

By substituting equation (10), we solve for free dCas9 concentration:

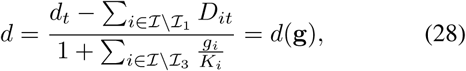

where g = [*g*_1_, *g*_2_,…, *g_n_*].

#### Senario (ii): Low Repression Level

When there are limited dCas9 protien or of gRNAs, most targets are not being repressed. Under this scenario, the number of free DNAs is approximately equal to the total number of DNAs: *D_i_* ≈ *D_it_*. Equation (22) can be written as:

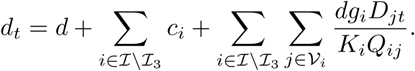

Next, we apply (10) to solve for free dCas9 concentration:

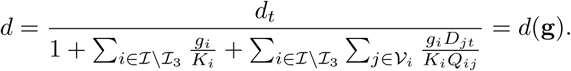

Finally, we focus on deriving the sign of *∂d*/*∂g_i_* for the most general cases.

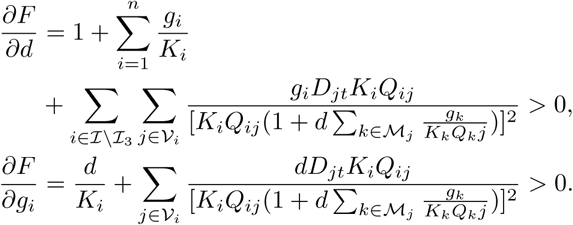

Since *∂F*/*∂d* ≠ 0 for all positive *d*, by the implicit function theorem, we have:

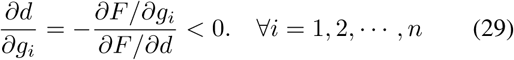

Consequently, the sign of the derivative *∂d*/*∂g_i_* is also guaranteed to be negative in a general network. This result will later be used to explore the sign of the *competition-induced hidden interaction* in section III-E.

### D. Summary

Since the dynamics of complexes *C_ij_* and *c_i_* have been set to QSS in all nodes, node dynamics can be reduced to that of the gRNAs and the protein. Therefore, from equations (19a), (20b), (21b) and the free DNA concentration obtained in (26), the dynamics in each type of node can be written as

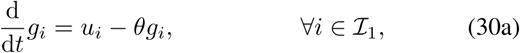

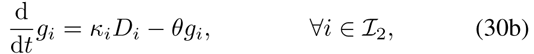

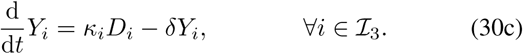

Using the results in (26), we can find the free DNA concentration *D_i_* to substitute into (30) to yield:

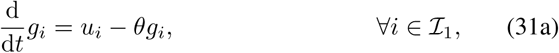

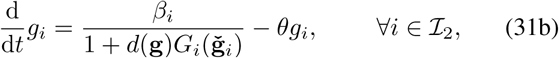

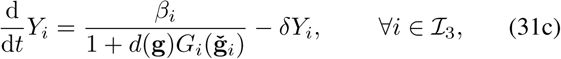

where we have defined

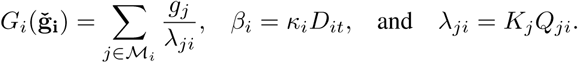

Equations (31b) and (31c) imply that the gRNA (protein) production in a type 2 (3) node is affected not only by its CRISPR regulatory inputs, characterized by 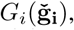 but also by the availability of dCas9 in the network *d*(g), which depends on the concentrations of all gRNAs in the network g.

**Fig. 5.**
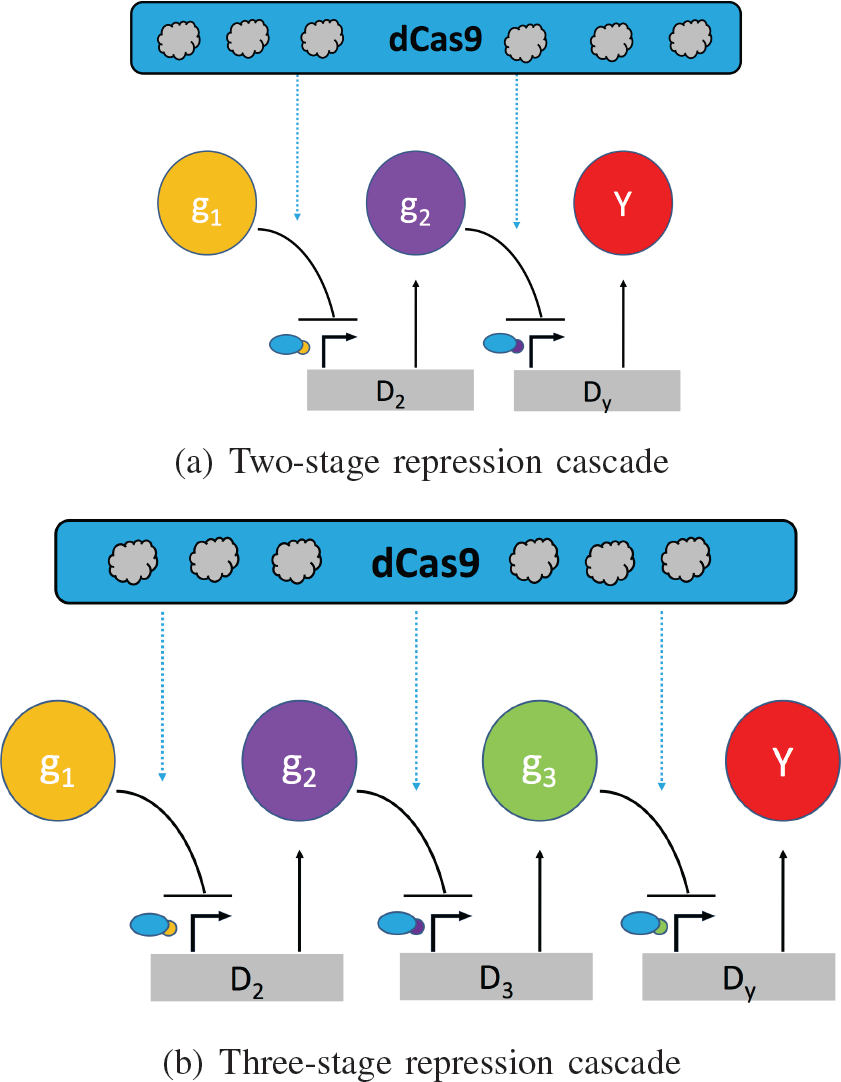
A simplified diagram for CRISPR-mediated repression cascades.

### E. Competition-Induced Hidden Interactions

Next, we address the question of how competition for limited dCas9 alters the intended interactions in a general network. To determine how dynamics of species x_*i*_ is affected by a change in the concentration of species x_*j*_, we introduce the notion of *competition-induced hidden interaction* from x_*j*_ to x_*i*_. In general, x_*i*_ can either represent a gRNA produced by a type 2 node 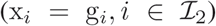 or a protein 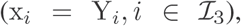 while x_*j*_ is a gRNA 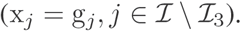

#### Definition

Consider the concentration dynamics of species x_*i*_:

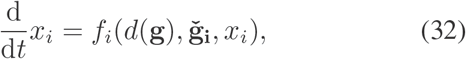

with *f_i_*(·) specified in (31b)-(31c). We define hidden interaction *E*(*x_i_*, *x_j_*) as:

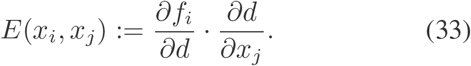

When *E*(*x_i_*, *x_j_*) = 0, we say there is no competition-induced hidden interaction from *x_j_* to *x_i_*. When *E*(*x_i_*, *x_j_*) > 0(< 0), there is a hidden activation (repression) on *x_i_* by *x_j_* due to resource competition. The hidden interactions can be graphically represented by a negative edge ⊣ from *x_j_* to *x_i_* if *E* < 0, and a positive edge ω if *E* > 0. The hidden interaction of cascades examples are drawn in Fig 6.

**Fig. 6.**
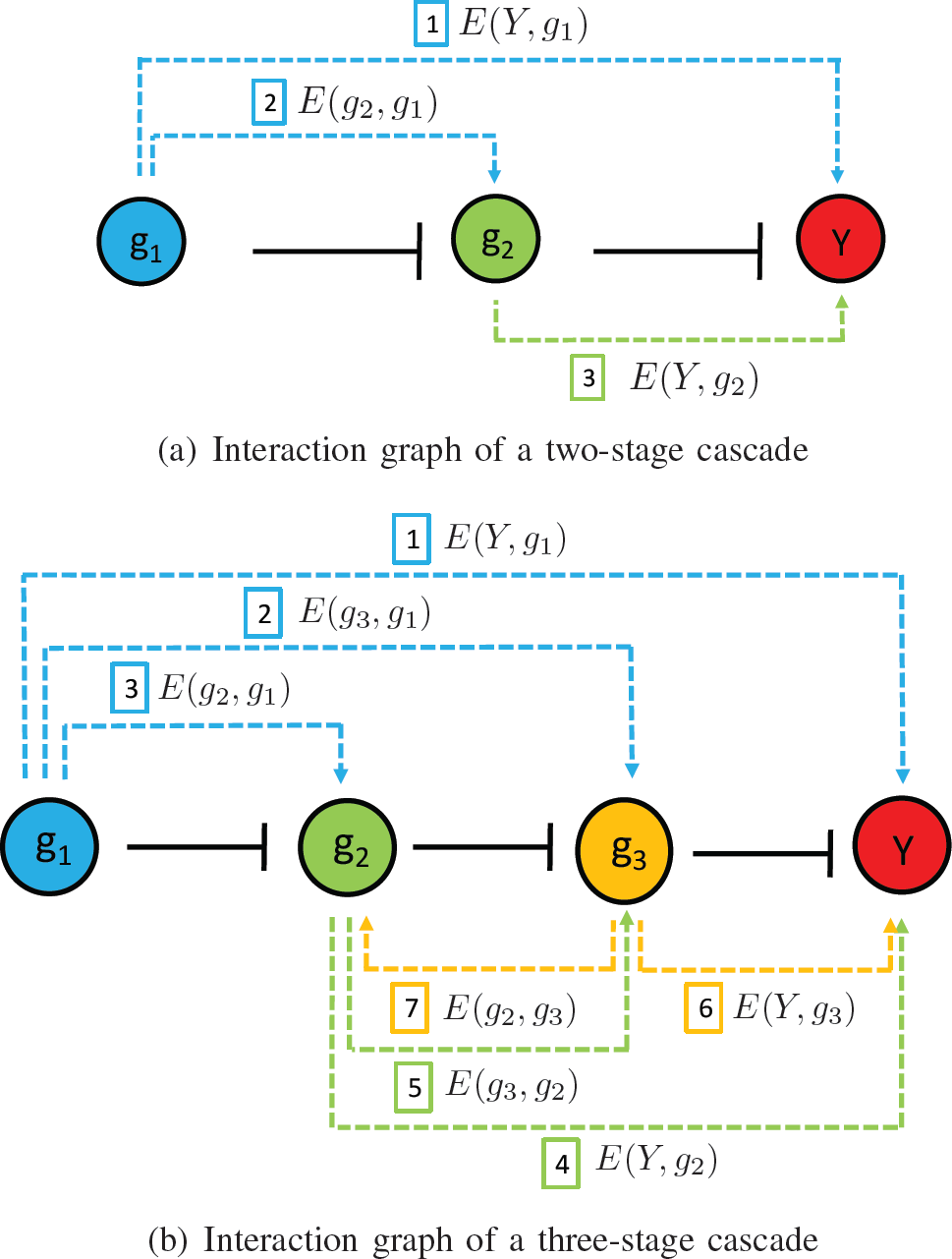
Interaction graphs of cascades: solid lines represent the original cascade structure; dashed lines represent the hidden interaction due to the resource competition effect.

#### Remark

All hidden interactions are activations for a repression network. This is because (i) *∂f_i_*/*∂d* < 0, as can be seen from (31), and that (ii) *∂d*/*∂x_j_* < 0 as we have shown in (29).

## IV AN EXAMPLE: REPRESSION CASCADES

### A. Two-Stage Cascade

Cascade circuits are one of the most common network motifs in both natural and synthetic gene networks. Here, we consider a simple two-stage repression cascade. The cascade is composed of three nodes: node 1, node 2 and node 3. In node 1, an external input *u*_1_ regulates the production of gRNA *g*_1_ 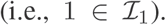. In node 2, the dCas9-gRNA complex *dg*_1_ regulates the production of gRNA *g*_2_ 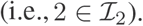 In node 3, the dCas9-gRNA complex *dg*_2_ regulates production of protein *Y* 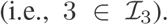 The structure of this motif can be represented by the interaction graph as g_1_ ⊣ g_2_ ⊣ Y. According to the general form (31a), (31b) and (31c), we have the following equations that describe the system dynamics:

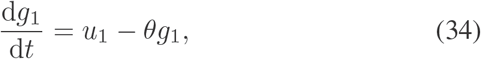

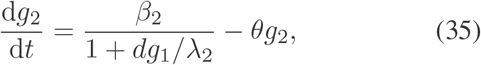

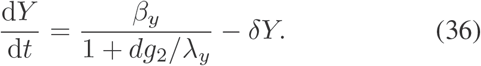

The hidden interactions resulting from the resource competition in illustrated in Fig.5(a). The steady state I/O response for the two-stage cascade is:

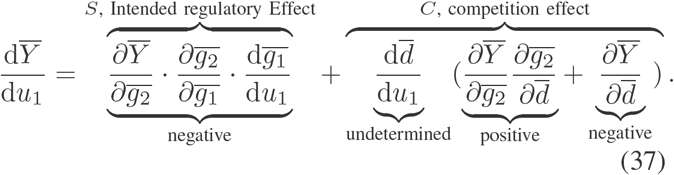

When there are abundant resources, the competition effect term vanishes *C* = 0 since free dCas9 concentration *d* remains approximately constant. It indicates that there is no resource competition effect in the system. However, when there are limited resources, free dCas9 concentration *d* fluctuates according to the gRNAs in the system *d* =*d* (g). Cmpetition effect *C* has to be considered. Since *sign* [*C*] is undetermined, the overall regulation from input to output is unknown in general. However, in this example, we demonstrate that perfect adaptation of output 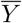 to resource fluctuation is possible by assuming *c_i_* ≫ *Q_i_*. To implement the assumption, one has to guarantee that there are more repressors formed than the targets in the system, such that after repressing the target, there are sufficient amount of free repressors left. From [20], we know that *Q_i_* ≈ 0.5nM, which makes the assumption easy to achieve. The assumption *c_i_* ≫ *Q_i_* is equivalent to *≫_i_* ≪ *dg_i_*, since *c_i_* = *dg_i_*/*K_i_*. Therefore, when *c_i_* ≫ *Q_i_*, the system dynamics is reduced to:

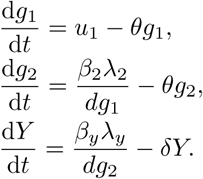

At steady state,

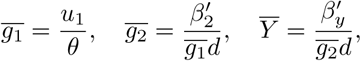

where 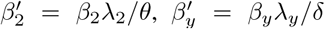 By calculating the partial derivatives of the steady state concentrations, the competition effect of the I/O response (37) is canceled out:

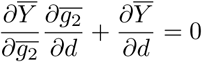

Therefore, we have the I/O response

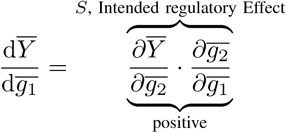

The competition effect *C* = 0 implies that perfect adaptation to fluctuations of the free resource *d* is achieved. In addition, the fact that the steady state output of the cascade is independent of *d* can be checked by substituting 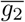 into 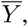, and the effective resource competition parameter *d* is cancelled out.

### B. Three-Stage Cascade

The three stage cascade is composed of four nodes: node 1, node 2 and node 4. The structure is similar to that of two-stage cascade with an addition node. In node 1, an external input *u*_1_ regulates the production of gRNA *g*_1_ 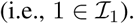 In node 2 (3), the dCas9-gRNA complex *dg*_1_(*dg*_2_) regulates the production of gRNA 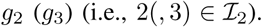 In node 4, the dCas9-gRNA complex *dg*_2_ regulates production of protein *Y* 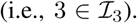 The structure of this motif can be represented by the interaction graph as g_1_ ⊣ g_2_ ⊣ g_3_ ⊣ Y. Like the two-stage cascade, the system dynamics can be derived from the general form (31a), (31b) and (31c):

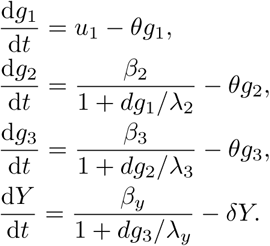

With the system dynamics, we draw the hidden interaction resulting from the resource competition in Fig.5(b). The steady state I/O response of a three-stage cascade is:

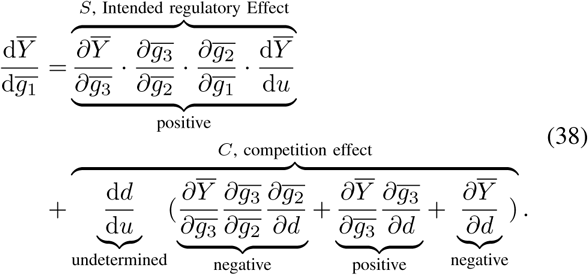

Since *sign* [*C*] is undetermined, the overall regulation from input to output is unknown. The hidden interaction resulting from the resource competition is in Fig.6(b). We apply the same assumption as the one we made for the two-stage cascade: *c_i_* ≫ *Q_i_*. Under this assumption, at steady state,

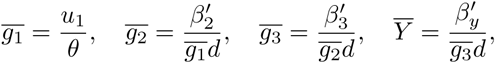

where 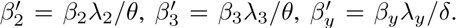 By the steady state concentrations, some of the terms in C of the I/O response (38) are cancelled out:

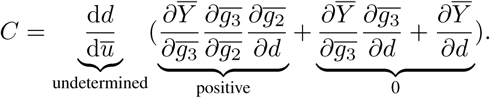

However, since there are remaining competition effects in the system (*C* ≠ 0)), the output concentration at steady state *Y* for a three-stage cascade is affected by resource competition. Fig. 7 shows how different amounts of resources affect the system behavior. In particular, when resources are abundant, the I/O behavior is as expected a monotonically decreasing function. As resources become limited, this behavior becomes biphasic.

**Fig. 7.**
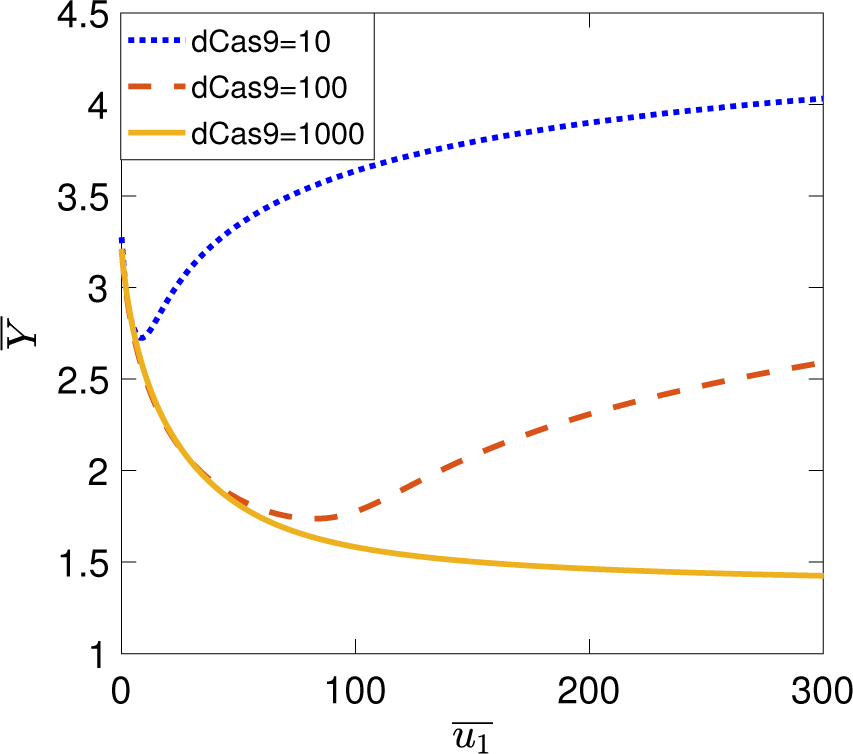
The yellow line shows that when there are sufficient resources, the steady state output monotonically decreases as the input is increase as expected from the regulatory interactions. The red line and blue line show that as *u*_1_ increases, the limiting resource results in an opposite effect on steady state output *Y*. It is noteworthy that the yellow line will eventually become biphasic if the input is further increased, which makes the resource become limited again. The parameters used here include *D_1t_* = *D_2t_* = *D_3t_* = 10nM, *K*_1_ = *K*_2_ = *K*_3_ = 0.01nM, *Q*_1_ = *Q*_2_ = *Q*_3_ = 0.5nM, δ= 1hr^−1^, = 100hr^−1^, *≫*_1_ = *≫*_2_ = 10hr^−1^, *≫*_3_ = 1000hr^−1^.

### C. n-Stage Cascade

The previous examples have demonstrated that resource limitation may affect the system to a different extent. For an n-stage cascade, it is composed of *n* + 1 nodes. In node 1, an external input *u*_1_ regulates the production of gRNA *g*_1_. In node *i*, where *i* = 2,…, *n*, the dCas9-gRNA complex *dg_i−1_* regulates the production of gRNA *g_i_* 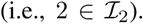 In node *n* + 1, the dCas9-gRNA complex *dg_n_* regulates production of protein *Y*. Under the assumption that *c_i_* ≫ *Q_i_*, where *i* = 1, 2,…, *n*, the dynamics of the system can be written as:

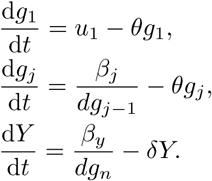

where *j* = 2, 3…, *n*. From the dynamics of repression cascades described above, for even-stage cascades, where *n* = 2*m*, and odd-stage cascades, where *n* = 2*m* + 1, the output concentrations at steady state are:

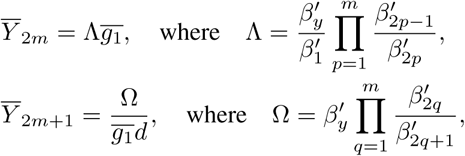

where 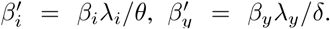 We conclude that perfect adaptation to resource fluctuations can be achieved in even-stage cascades since 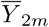 are independent of the resource concentration (*d*). Odd-stage cascades, on the other hand, have the output concentration at steady state 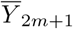 depending on *d*. Therefore, when designing cascades in CRISPR-based regulation networks, even-stage cascade may be preferable to odd-stage cascades.

## V. DISCUSSION AND CONCLUSIONS

CRISPR-based transcription factors provide an exciting alternative to synthetic TF design due to their ease of use and efficiency in regulating multiple genes in parallel. With CRISPR regulators, multiple endogenous genes can even be regulated to control a complex phenotype, such as cell fate [23]. As the complexity of CRISPR-mediated gene regulation networks increases, it is critical to identify the underlying resource-sharing mechanism. In this paper, we first develop a simpler modeling framework: the parallel model. We trained the parameters in the model with preliminary experimental results. Our simulation results showed that the competition effect can cause around five folds of hidden activation, which matches the experimental results. Then we proposed a general modeling framework which is applicable to all repression networks. Finally, we use repression cascades as an example to illustrate that circuits in certain configurations possess a better robustness to the competition effect, which can alter the sign of steady state I/O response. In the future, we will examine how CRISPR-mediated gene regulation networks can be designed to be more robust to resource competition effects. The outcome of the examination will then be validated with further experiments.

## Acknowledgement

We thank Aaron Dy, Massimo Bellato and Cameron McBride for helpful discussions.

